# VIRUSBreakend: Viral Integration Recognition Using Single Breakends

**DOI:** 10.1101/2020.12.09.418731

**Authors:** Daniel L. Cameron, Anthony T. Papenfuss

**Author notes:** To whom correspondence should be addressed: Daniel L Cameron and Anthony T. Papenfuss.

## Abstract

Integration of viruses into infected host cell DNA can causes DNA damage and can disrupt genes. Recent cost reductions and growth of whole genome sequencing has produced a wealth of data in which viral presence and integration detection is possible. While key research and clinically relevant insights can be uncovered, existing software has not achieved widespread adoption, limited in part due to high computational costs, the inability to detect a wide range of viruses, as well as precision and sensitivity. Here, we describe VIRUSBreakend, a high-speed tool that identifies viral DNA presence and genomic integration recognition tool using single breakend variant calling. Single breakends are breakpoints in which only one side has been unambiguously placed. We show that by using a novel virus-centric single breakend variant calling and assembly approach, viral integrations can be identified with high sensitivity and a near-zero false discovery rate, even when integrated in regions of the host genome with low mappability, such as centromeres and telomeres that cannot be reliably called by existing tools. Applying VIRUSBreakend to a large metastatic cancer cohort, we demonstrate that it can reliably detect clinically relevant viral presence and integration including HPV, HBV, MCPyV, EBV, and HHV-8.

## Background

As made abundantly clear by the SARS-COV-2 and HIV pandemics, viral infections constitute a major worldwide threat to human health. While most viruses do not integrate into the host genome, there is a significant global health burden caused by the subset of those that do, especially in cancer ^1^. For example, human papillomavirus (HPV) is present in the majority of cervical cancers, Merkel cell polyomavirus (MCPyV) is the primary cause of Merkel cell carcinoma, and the Epstein-Barr virus (EBV) infects around 90% of the human population is associated with multiple forms of cancer ^2^. Other oncoviruses include Kaposi’s Sarcoma-associated herpesvirus (HHV-8), and Hepatitis B virus (HBV)—the leading cause of Hepatocellular carcinoma (HCC). For some of these, the location of the viral integration is a direct driver of oncogenesis with HBV integrations in the TERT promoter region are associated with high telomerase expression and cancer cell survival ^3^. This integration site-specific behaviour is not just limited to oncogenic viruses, as human immunodeficiency virus (HIV) elite controllers have shown to have a high rate of centromeric viral integrations^4^. The reliable detection of viral integrations anywhere in the genome is key to understanding the effect of viral integration to disease.

Recent advances in sequencing technology have made routine large-scale whole genome sequencing (WGS) possible, including tumour sequencing ^5^. These WGS data sets enable the detection of viral integrations through the identification of structural variant breakpoints between the host genome and the viral sequence. While there exist several tools capable of detecting viral integrations in WGS data, these tools have not yet gained widespread adoption. Existing tools fall short in one or more of three areas: the ability to detect more than one virus or virus family, runtime performance, and the inability to detect integrations into repetitive regions of the host genome (such as centromeres).

At a high level, WGS viral integration detection software finds integration sites by identifying clusters of reads or read pairs spanning from host reference sequence to viral reference sequence. Viral integration tools such as BatVI ^6^, VirTect ^7^, and Virus-Clip ^8^ require a viral reference as input. While some of these tools are true single-virus tools, others can in theory be configured with multiple viral reference genomes. Including related viruses causes read alignment ambiguities when these viruses contain homologous regions. VirusFinder ^9^, VirusFinder2/VERSE ^10^, and VirusSeq ^11^ avoid this problem by first identifying viral presence before proceeding to integration detection using a single viral reference genome. These tools are still limited to a single viral reference genome, so a HHV-6 infection may mask the presence of a short genome such as HBV. The Pan-Cancer Analysis of Whole Genomes (PCAWG) project ^3^ avoided this problem by performing viral read classification prior to viral integration detection but this pipeline is not generally available as a standalone tool and its integration detection performance is determined by their use of VERSE as the sole integration detection tool. The need for an integrated, easy to use, virome-wide integration detection was identified by Chen at al^12^, a gap we propose to fill with VIRUSBreakend.

For the vast majority of whole genome sequencing projects, viral integration detection is only one part of a larger analysis. Tools that are not computationally efficient will struggle to gain widespread adoption. Tools such as BatVI and Virus-Clip were developed in direct response to the computational cost of tools such as VirusFinder, VERSE, and VirusSeq. By far, the most computationally expensive step is the alignment of reads to the host and viral genomes and multiple approaches have been taken. VERSE, VirusSeq, and ViralFusionSeq ^13^ use a host then virus alignment approach, BATVI and Virus-Clip use a virus then host approach, while ViFi ^14^ and VirTect ^7^ take a combined host and virus approach. Each of these approaches have their advantages and drawbacks, but host then virus alignment approach has the unique advantage that it uses as input a bam file that will almost certainly have been generated in a typical WGS pipeline. Here, we show that this approach reduces the real-world computational cost of incorporating viral integration detection is less than one tenth of the computational cost of realignment.

Finally, and most crucially, all existing viral integration detection tools rely on clusters of read alignments. Even tools such as VirusFinder/VERSE that perform an assembly step, still relying on read host alignment clusters to determine the insertion site. This fundamentally limits the viral integration detection capability in host regions with low mappability. Ignoring low mappability reads will result in false negatives, and reporting either a single arbitrary alignment or all possible alignments of a multi-mapping reads with both result in a high false positive rate and overestimation of the number of insertion sites. Our solution to this problem is to perform single breakend variant calling on the viral reference genome. Single breakends are breakpoints in which only one side is uniquely aligned to the reference genome ^15^. By first identifying where in the viral genome an integration site occurs and assembling the host sequence adjacent to the integration, we obviate the problem of multi-mapping host read alignments. In cases where the host location cannot be unambiguously determined from the assembled contig, the contig sequence provides information about the repeat context of the integration site.

Here we present a novel single breakend-based approach that can reliably detect viral integrations anywhere in the host genome. By identifying and assembling single breakend variants in the virus genome followed by taxonomic classification and alignment of the breakend contigs, VIRUSBreakend is able to reliably identify viral integrations in regions inaccessible to current integration detection approaches.

## Results

### VIRUSBreakend Overview

VIRUSBreakend uses a multistage approach to identifying viral insertions (Figure 1). Starting with a host-aligned SAM/BAM/CRAM file, VIRUSBreakend identifies viral reads of interest through Kraken2 ^16^ taxonomic classification of all unaligned or partially aligned sequences using. If the read is at least partially classified as a virus, the full read pair is considered for further analysis.

**Figure 1:**
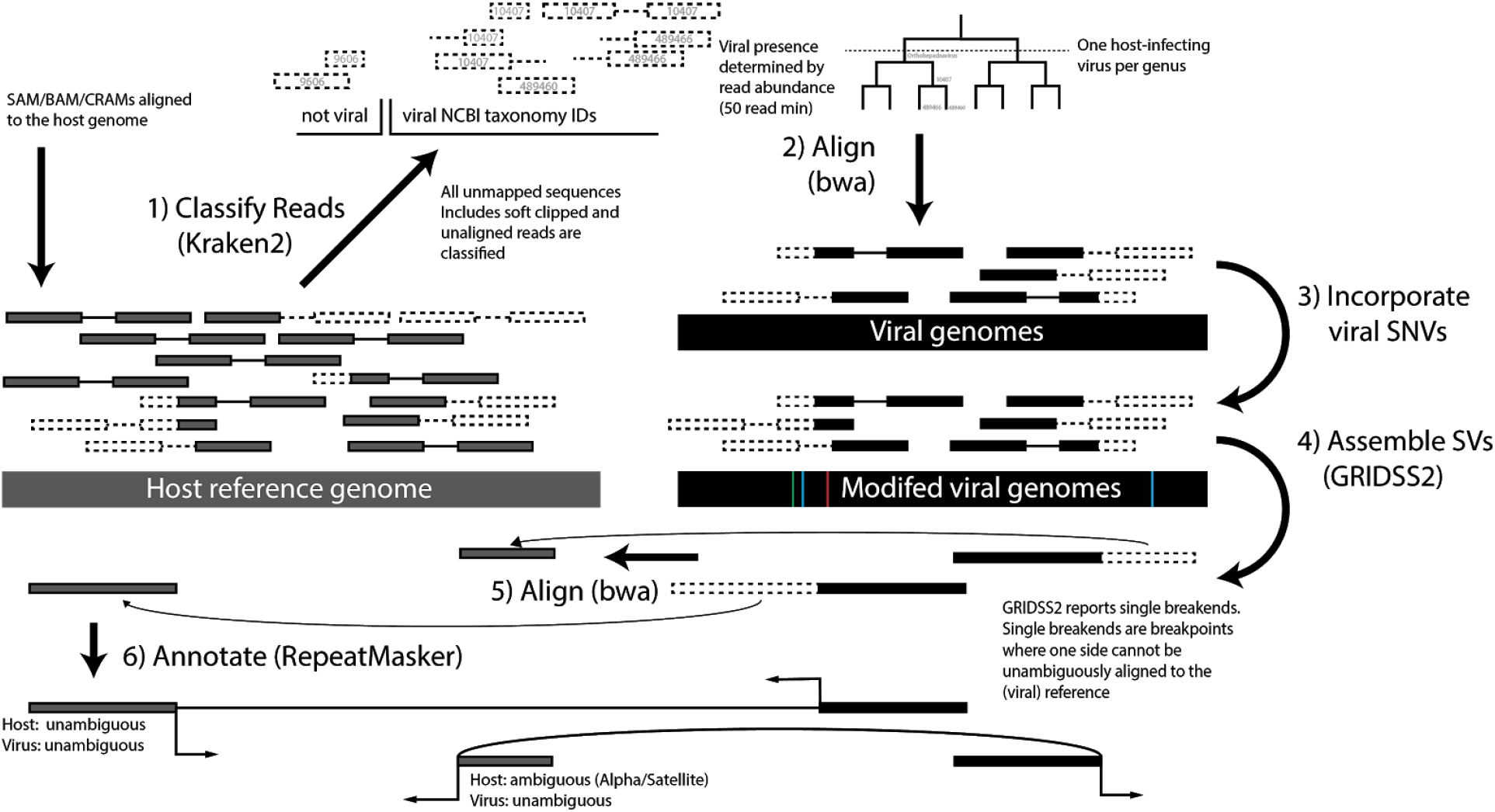
VIRUSBreakend workflow. Sequences not aligned to the host are taxonomically classified to identify viral abundance. The most abundant host-infecting virus per genus is incorporated into a viral reference. The full read pairs for all viral sequences are aligned, SNVs are called and incorporated into the viral reference. Single breakends variants are assembled and called and host integration sites identified by alignment of the breakend assemblies. This virus-centric approach allows identification of integrations in repetitive/low mappability host regions.

Viral read pairs are aligned to a viral reference consisting of the most abundant human-infecting virus in each genus. SNVs called with bcftools and the viral reference modified to incorporate these SNVs. Viral read pairs are realigned to the updated reference and structural variants called using GRIDSS2 ^17^ and filtered to single breakends. Single breakends are breakpoints in which one side cannot be unambiguously aligned to the (viral) reference. In the case of viral integration breakpoints, this is because the viral reference does not include the host genome. As well as the location and orientation of the break junction, GRIDSS2 reports the assembled sequence for every single breakend.

The assembled single breakend sequence is used to identify the host integration site by aligning to the host reference. Identified breakpoints fall into two categories: sites in which the host mapping is unambiguous, and ambiguous sites in which the integration site cannot be unambiguously determined (such as integrations into alpha satellite repeats). To facilitate downstream analysis, integration sites are annotated with the RepeatMasker ^18^ repeat type and class of the single breakend sequence.

### Synthetic Benchmark

To evaluate theoretical performance, we created a synthetic benchmark with realistic insertion sites and compared VIRUSBreakend to VERSE, ViFi, and BATVI, as well as GRIDSS2. GRIDSS2 was included both as a breakpoint caller against a reference containing the human and viral genomes, as well as to evaluate the performance of host-centric single breakend calling, and was selected due to its performance ^19,20^. Insertion sites were simulated by inserting 2000bp of the non-reference HBV strain LC500247.1 at each 1Mb interval along chromosome 1 of the Telomere-to-Telomere consortium whole genome assembly of CHM13. Viral integration callers were run using their default settings with hg19 used as the host reference genome. To account for differences between CHM13 and hg19 coordinates, insertions were considered correct and counted as true positives if the viral coordinates were within 750bp and the insertion site coordinates were within 1Mb. Insertions were considered homologous calls if matching only on the viral side.

Using the F-score to evaluate performance, VIRUSBreakend outperforms the specialised callers at 30x coverage with an F-score of 0.985 compared to 0.84 for BATVI, 0.9 for VERSE, and 0.82 for ViFi. A similar result is observed at 10x, 15x and 60x, where VIRUSBreakend achieves F-scores of 0.90, 0.94 and 0.997 respectively compared to 0.81, 0.85 and 0.82 for BATVI, 0.63, 0.90, and 0.93 for VERSE, and 0.77, 0.82 and 0.83 for ViFi. At 5x coverage, VIRUSBreakend comes in second by a small margin (F-score 0.66) and is outperformed by BATVI (F-score 0.67) compared to 0.01 (VERSE) and 0.45 (ViFi). GRIDSS2 host-centric single breakend variant calling does not perform particularly well (F-scores: 0.71, 0.80, 0.82, 0.83 and 0.84 at 5x, 10x, 15x, 30x and 60x), but breakpoint calling using a combined host and viral reference has the best performance at low coverage (F-scores: 0.91, 0.94, 0.93, 0.90, 0.87 at 5x, 10x, 15x, 30x and 60x). BATVI and GRIDSS2 breakpoint calling show almost identical false positive rates that increase with coverage whereas ViFi shows the opposite trend with a higher false positive rate at low coverage. ViFi, VIRUSBreakend and GRIDSS2 single breakends all have negligible false positive rates in this simulation. Only 4 of insertion sites were uniquely called by VIRUSBreakend indicating that, collectively, existing callers are able to identify most integration sites, but not reliably so.

### Hepatocellular Carcinoma Benchmark

Next, we evaluated sensitivity on the 22 PCR/Sanger validated hepatitis B virus (HBV) integration sites from a hepatocellular carcinoma cohort^21^ used by VERSE ^10^ and ViFi ^14^ VERSE identified 13 integration sites, ViFi 12, GRIDSS2 15, BATVI 16, and VIRUSBreakend 15. 4 of the validated integrations were entirely within HSATII or Beta satellite repeats for which VIRUSBreakend reported integrations into HSATII/Beta satellite repeats with a higher sequence similarity to the assembled viral integration sequence than the nominal integration site. Since the host PCR primer sequences chosen are present at both at the validated sites and the sites called by VIRUSBreakend, it is unclear where the actual integrations occurred. The locations are homologous and integration in either location would result in successful PCR amplification. Treating these homologous sites as correct calls, the sensitivity of VIRUSBreakend rises to 19/22. The remaining three VIRUSBreakend false negatives were due to insufficient coverage for assembly of the reads supporting the insertion site to be successful.

### Hartwig Medical Foundation Cohort

To evaluate performance on a large cohort, we ran VIRUSBreakend on 5,191 tumour samples from the Hartwig Medical Foundation metastatic tumour cohort ^5^. We detected viral presence in 610 samples, with integration in 160. The most prevalent viruses were EBV with viral integration detected in 27 of 278 samples with viral presence, and HHV-6 (33/144), HPV-16 (60/93), HHV-7 (3/25), and HPV-18 (21/21), and HHV-5 (0/19). The likelihood of integration detection was driven primarily by viral coverage with at least one integration site found in 90% (121/135) of samples achieving 10x viral coverage (Figure 3a).

Of the 43 cervical cancer in the cohort, 38 were HPV positive, with integration sites found for 37 (Figure 3b) with anal (11/20), penile (6/11), oropharynx (4/10), cancers also enriched for HPV. As expected^22^, the 5 HPV negative cervical cancers were the only cervical cancers with TP53 driver mutations. were also enriched for HPV. Of the 9 HBV+ samples, 7 were in liver cancers (n=19) (all with detected integrations), which is consistent with previous findings ^3^. Recurrent HBV integration was found in TERT (4 samples), and likely driver integrations found in/upstream of FOXP2, WNT2, EML6, ZDHHC11, and CTSC. Merkel cell polyomavirus integration was detected in all 6 Merkel cell carcinomas. HHV-8 was detected in the single Kaposi’s sarcoma sample in the cohort, although the integration site was not.

Whether EBV is a risk factor for lung cancer is still subject to debate ^23^. While we did find EBV viral presence enriched in lung cancer (69/666, p=0.000002), only 2 integration sites were found. In all samples, viral depth of coverage was less than 2.5% of the host indicating that EBV is not clonally integrated into the tumour in any sample.

Integration of viral sequence into unmappable regions of the genome was dominated by herpesvirus telomeric integration as expected ^24^. While recently developed optical mapping protocols are able to localise these viral integrations to specific chromosomes for inherited chromosomally integrated HHV ^25^, VIRUSBreakend can detect telomeric integrations but lacks the long range information required for disambiguation. The unmappable HPV integrations predominantly occurred in samples in which a mappable HPV was also found indicating that these may be passenger events. Only 3 HPV-16 samples contained only unmappable HPV integrations: a possibly intronic poly GGAA; an alpha satellite integration; and a highly amplified telomeric integration. Further investigation will be required to ascertain the functional relevance of centromeric and telomeric viral integrations.

### Runtime performance

To evaluate runtime performance, all callers were run on the HCC 177T sample with 4 threads specified (Figure 2c). Each caller was allocated 4 cores and 20GB of memory on a HPC cluster containing dual core Xeon E5-2690 servers. Both VIRUSBreakend and VERSE were using host-aligned BAM as inputs, whereas BATVI and ViFi used fastq input. VIRUSBreakend completed in 35min, BATVI 41 hours, VERSE 17 hours, and ViFi 44 hours.

**Figure 2:**
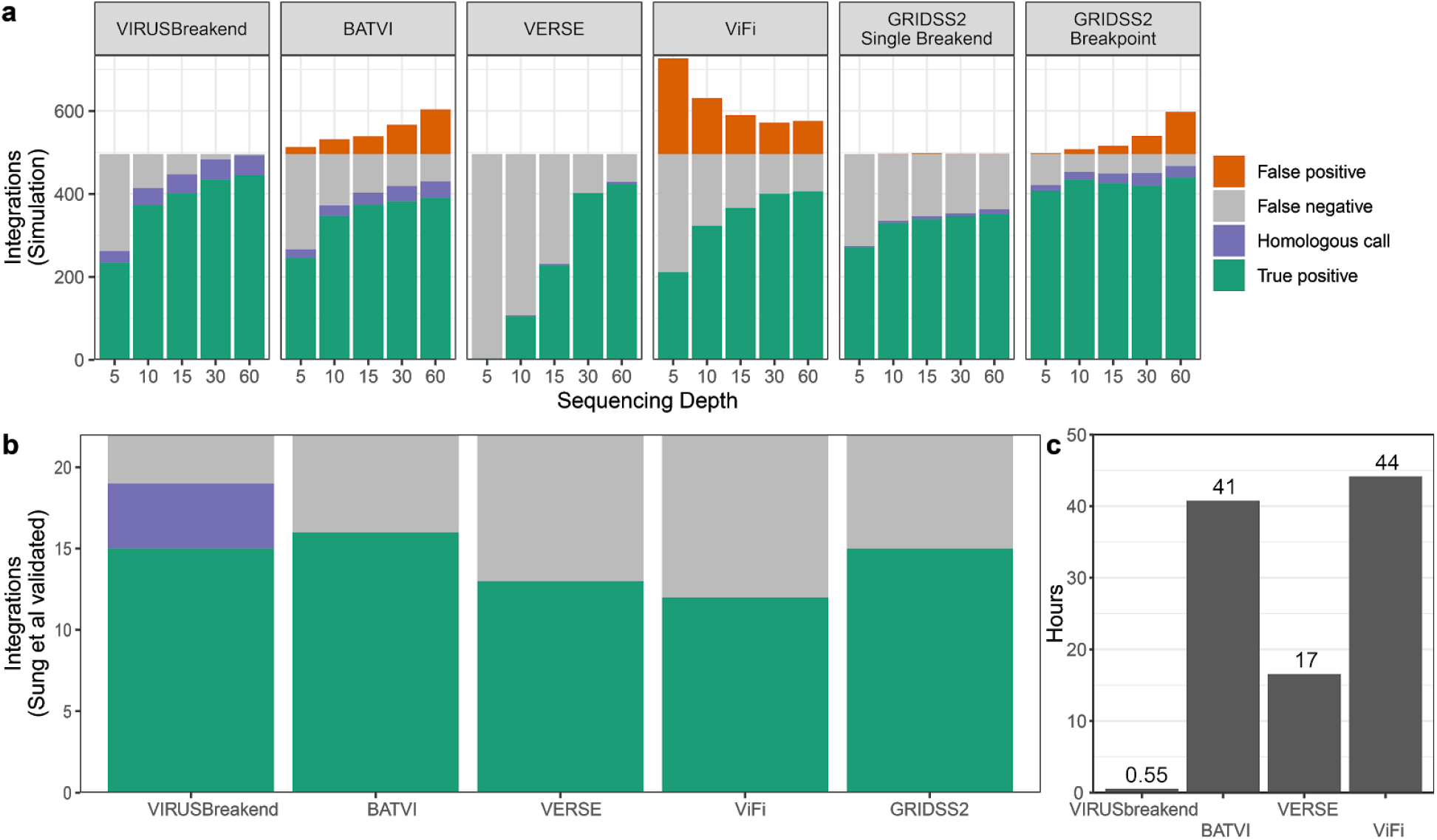
a) Performance of simulated HBV integrations. 2kbp of the LC500247.1 HBV strain were integrated at every Mb of chromosome 1 of the complete CHM13 reference genome for a total of 248 insertion sites. Callers were run against hg19. Calls were considered homologous true positives if the viral position matched but the integration site was at a homologous location in the human reference. GRIDSS2 single breakend calls are against a host-only reference, and GRIDSS2 breakpoint calls are against a combined host and viral reference. On this data set, VIRUSBreakend achieves perfect precision and outperforms (f-score) all other callers above 10x coverage. b) Sensitivity on the 22 integration sites validated by Sung et al 2012 in a hepatocellular carcinoma cohort. The 3 VIRUSBreakend false negatives had insufficient coverage for the integration site to be assembled. c) Runtime on sample 177T from the hepatocellular carcinoma cohort when allocated 4 cores and 20GB memory. The cost of the initial host genome alignment is not included for VIRUSBreakend or VERSE as this is typically performed in a WGS pipeline regardless of whether viral integration detection is performed or not.

**Figure 3:**
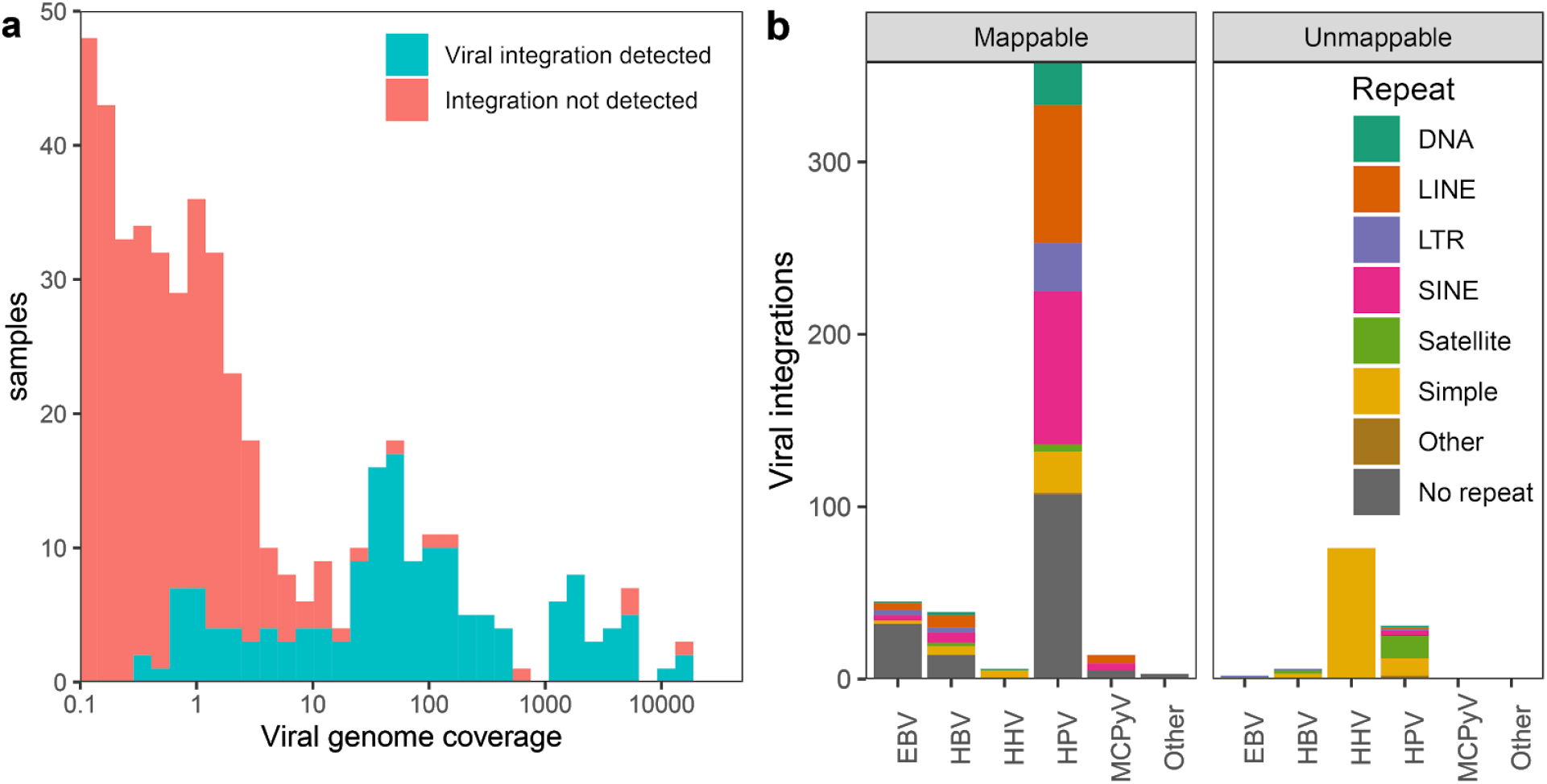
Analysis of 5,191 metastatic tumour samples from the Hartwig Medical Foundation cohort. a) Distribution of viral genome coverage. Viral integration sites were not found for samples with low viral coverage. b) Mappability of integration sites. Unmappable sites have multiple candidate integration sites and are dominated by HHV integration into telomeric repeats and HPV integration

On the Hartwig Medical Foundation cohort, VIRUSBreakend average execution time on a 4 core 16Gb c2-standard-4 google cloud compute instance was 45 minutes for samples without detected virus and 85 minutes for samples with detected virus. VIRUSBreakend runtime is dominated by input BAM/CRAM decompression with CRAM decompression requiring more CPU usage than BAM. VIRUSBreakend supports direct streaming of input files. This eliminates the input file copy overhead when run in the cloud. Using input file streaming and preemptible instance c2-standard-4 instances, the entire Hartwig cohort of over 500TB of CRAMs was processed for under US$500.

## Discussion

When viruses are integrated into low mappability sequences, existing read mapped based approaches must choose between erring on the side of caution and omitting these calls, or aiming for high sensitivity at the cost of a high false discovery rate. By taking a virus-centric single breakend approach, VIRUSBreakend solves this dilemma and enables both accurate and sensitive integration detection even in regions of low mappability. This does however come at a cost. Since the single breakends must be assembled, a traditional read mapping based caller will have greater sensitivity on low coverage samples as they do not suffer from the abrupt drop in sensitivity that VIRUSBreakead is subject to when there is insufficient coverage for reliable assembly. Similarly, above ~3000x viral genome coverage, the assembler used by VIRUSBreakend starts hitting assembly graph complexity limits thus making integration site detection somewhat unreliable for very highly expressed DNA viruses. Key to the extremely fast runtime performance of VIRUSBreakend is use of host-aligned input enabling the vast majority of reads to be immediately discarded. If a host-aligned input file was not available, the computational cost of VIRUSBreakend would increase by over an order of magnitude. For the vast majority of projects, this requirement is unproblematic as a host-aligned BAM/CRAM will be created for variant calling purposes. Since VIRUSBreakend has no constraints on the host reference used (other than they not contain the viral sequences), it is suitable for incorporation into an existing WGS pipeline.

Finally, while the approach of Kraken2 taxonomic classification to identify the viral reference genomes does allow pan-virome integration detection, it does come with limitations. The one genome per genus ensures that a small amount of taxonomic misclassification does not result in a viral reference containing multiple related viral strains, but it also masks the presence of genuine viral co-infection by closely related viruses. Similarly, since the viral database contains only the exemplar RefSeq reference for each taxonomic identifier, distantly related strains may not be identified at all. This could be mitigated through the use of a more comprehensive viral database.

## Conclusions

The single breakend variant calling and assembly approach taken by VIRUSBreakend enables sensitive and accurate viral integration detection even in low mappability host regions. Since it combines both viral presence and integration detection into a streamlined high-speed tool, it is ideal for augmenting WGS-based sequencing pipelines with viral information and has direct clinical utility for WGS cancer patient reporting. VIRUSBreakend is a marked improvement on existing tools and provides a foundation for future research into the impact of viral integration into centromeric and telomeric regions currently considered inaccessible to short read sequencing.

## Methods

### VIRUSBreakend pipeline

VIRUSBreakend uses a multistage approach to identifying viral integration sites. As input, it uses a SAM/BAM/CRAM file of reads aligned to the host reference genome. Viral reads are classified using Kraken2 ^16^, aligned to the most abundant host-infecting virus with bwa ^26^, realigned to a modified viral reference that incorporates SNVs called by bcftools, single breakends identified with GRIDSS2 ^17,27^, aligned to the host to identify putative integration sites, and annotated with RepeatMasker to identify false positive and multi-mapping integration sites.

All read sequences 20bp or longer that are not aligned to the host reference genome are classified using a Kraken2 database containing the human, viral and UniVec_Core sequences. For soft clipped reads, only the unaligned bases are classified. For split read alignments, only the bases not aligned to either location are classified. The full read is considered for all unmapped reads.

Sequences are considered to be of interest if either Kraken2 classifies the overall sequence with a viral taxid, any kmer is classified with a viral taxid and all kmers between that kmer are an ancestor of a viral taxid. That is, the sequence is entirely a virus of interest, or could be a split read containing a virus of interest and non-viral sequence (such as a read overlapping a host integration site).

The originating reads names for sequences of interest are tracked and a second pass over the input file is performed to extract the entire originating fragment (i.e. both reads if paired-end sequencing) for all sequences of interest. A viral reference is created consisting of the most abundant human-infecting viral taxid for each genus. Here, we define ‘most abundant’ as a viral taxid of interest with the most reads directly assigned to that taxid by Kraken2, with ties broken by selecting the taxid with the most read assigned to that taxid or any of the descendent taxid. Only taxid with at least 50 sequences to it or a descendant and having an associated genome sequence in the kraken2 database are considered. If no taxid of interest reaches the 50 sequence threshold, processing terminates. In the case of multiple genomes associated with a taxid of interest, only the first contig is incorporated into the viral reference. Any accession included in the NCBI Viral Genomes ^28^ neighbours file (https://www.ncbi.nlm.nih.gov/genomes/GenomesGroup.cgi?taxid=10239&cmd=download2) with host containing “human” is considered human-infecting.

Extracted reads are aligned to the viral reference and SNVs called using bcftools call -c -v --ploidy 1 -V indels. The viral reference genome is then updated using bcftools consensus and extracted reads realigned to the new reference. To preserve viral coordinates, indels and SVs are not incorporated. Assembly/structural variant calling is performed using GRIDSS2. To improve library fragment size distribution estimation and GRIDSS quality score calculations, GRIDSS metrics are precomputed from the first 10,000,000 reads in the host-aligned input SAM/BAM/CRAM.

Candidate host genome integration positions are identified by aligning the breakend inserted sequenced to host reference genome using bwa/gridss.AnnotateInsertedSequence which annotates variants with the nominal alignment, mapq, and also any alternative alignments reported in the bwa XA tag. The GRIDSS VCF is annotated with gridss_annotate_vcf_repeatmasker.sh and gridss_annotate_vcf_kraken2.sh using the same custom Kraken2 database used for viral sequence identification.

The VCF is filtered to only single breakend variants in which the kraken2 classification of the single breakend sequence matches the host NCBI taxonomy ID (human, 9606). Finally, single breakend calls are transformed into host-virus breakpoint calls based on the first alignment position reported by bwa/gridss.AnnotateInsertedSequence with the alignment mapping quality score used to determine whether the called position is ambiguous or not.

### Tools

VIRUSBreakend 2.10.2, BATVI 1.03, VERSE 2.0, ViFi (commit d56f4c2) and GRIDSS 2.10.2 were run with default settings using the installation procedures outlined in their user guide. All VERSE perl scripts were edit to use “#!/bin/usr/env perl” so as to be compatible with a conda installation, the base qual cutoff for the embedded CREST was reduced to 15 since wgsim defaults to 17 as a base quality score, and the undocumented perl dependencies were iteratively installed until execution no longer raised missing library errors. The missing step of creating bwa index in the BATVI batmis directory was run in addition to the installation instructions to circumvent the fatal error encountered building the indexes when following the documented instructions. BATVI full call set results (predictions.opt.subopt.txt) were not included as the recall of this call set was lower than the BATVI high confidence calls on the simulation data. The following modifications were required to get the most recent version (27 Mar 2019 d56f4c28) of ViFi to run: recommendation to use the supplied docker image was ignored since the docker command-line parsing logic was incorrect and crashed when using -b, and ignored to -v parameter thus always ran against HPV; crash bug in scripts/get_trans_new.py:238 was fixed by correcting the incorrect parenthesis on line 233; a conda environment was created using “conda create -n vifi bwa=0.7.17 python=2.7 pysam=0.15.2 samtools=1.9 hmmer”. VIRUSBreakend was run using GRIDSS version 2.10.2, 4 threads and --rmargs “-e rmblast” since the conda installation of RepeatMasker does not perform RepeatMasker configuration. RepeatMasker 4.1.0 and Kraken 2.1.0 were installed from BioConda ^29^. Reads were aligned to hg19 using bwa mem 0.7 and converted to bam and coordinate sorted using samtools 1.11.

### HCC Benchmark

Reads associated with samples 145T, 177T, 180N, 186T, 198T, 26T, 200T, 268T, 43T, 46T, 70T, 71T, 95T in project ERP001196 were downloaded from SRA using fasterq-dump from sra-tools 2.10.8. ERR093473 and ERR173541 were excluded from analysis due non-transient fasterq-dump errors that could not be rectified. Since VIRUSBreakend supports multiple input files and GRIDSS performs per-file library fragment size distribution estimation, bam files were not merged. GRIDSS calls were annotated with gridss_annotate_kraken2.sh and filtered to single breakends with a viral taxid. VERSE and ViFi were not rerun and the results presented in their respective publications were taken as is.

### Simulation

Simulated reads were generated using targeted insertion sequences between version of the 1.0 chr1 telomere-to-telomere consortium assembly of chm13 (https://github.com/nanopore-wgs-consortium/CHM13) and LC500247.1. Each targeted inserted sequence was processed and called independently with insertions spaced evenly at every Mb of chrq chm13. 50kbp of chm13 from 1,000,000n - 50,000, 1,000,000 was concatenated with 1kbp of LC500247.1 sequence from 4n to 4n + 2,000 which was then concatenated with 50kbp of chm13 from 1,000,000n + 10, 1,000,000 + 50,010. This resulted in an insertion of 2kbp of HBV, a 10bp gap between the left and right side of the integration, and 50kbp of human sequence flanking the insertion site. GRIDSS metrics from 20M simulating reads from chm13 using the same parameters were used to emulate realistic VIRUSBreakend WGS metrics calculations as no simulation data set contained the 10M reads used for metrics approximation. Read were simulated using ART 2.5.8 ^30^ with parameters --noALN --paired --seqSys HSXn -ir 0 -ir2 0 -dr 0 -dr2 0 -k 0 -l 150 -m 500 -s 100 -rs 1. Separate data sets were generated with coverage (--fcov) of 5, 10, 15, 30, and 60. An E. Coli read pair and an E Coli/human read pair was appended to each fastq file to ensure ViFi did not crash. VIRUSBreakend was run with --minreads 15 to ensure the viral presence filtering did to interfere with this benchmark of integration detection capability. GRIDSS2 calls were filtered to single breakend variants that realigned to HBV. To account for coordinate differences between hg19 and CHM13 and LC500247.1 and the HBV references used by the callers, calls were considered a true positive if the hg19 host position was within 1Mb of the CHM13 truth position, and the viral position was within 750bp of the LC500247.1. Calls were considered homologous calls if the viral position matched and no full matches were found for that breakpoint. Duplicate or unmatched calls were considered false positives. GRIDSS2 breakpoint calls were generated by concatenating NC_003977.2 to hg19 aligning using bwa mem 0.7.17 and filtering to breakpoint calls involving NC_003977.2.

### Hartwig Medical Foundation

VIRUSBreakend was run on 1988 samples in the Hartwig Medical Foundation cohort using pre-emptible c2-standard-4 (4 vCPUs, 16 GB memory) instances. --gridssargs “--jvmheap 13g” was specified since the GRIDSS2 JVM defaults to a 30GB heap size. TTV and xenotropic retroviruses were excluded from counts.

## Availability of data and materials

VIRUSBreakend is available as free and open source software under a GPLv3 license and is available at http://github.com/PapenfussLab/gridss/

Hartwig Medical Foundation cohort data was obtained from the Hartwig Medical Foundation (Data request DR-005). Standardized procedures and request forms for access to this data can be found at https://www.hartwigmedicalfoundation.nl/en.

## Competing interests

The authors declare no competing interests.

## Funding

A.T.P. was supported by a National Health and Medical Research Council (NHMRC) Senior Research Fellowship (1116955) and the Lorenzo and Pamela Galli Charitable Trust. D.L.C. and A.T.P. were supported by an NHMRC Ideas Grant (1188098). The research benefitted by support from the Victorian State Government Operational Infrastructure Support and Australian Government NHMRC Independent Research Institute Infrastructure Support.

## Authors’ contributions

DLC designed and implemented VIRUSBreakend. DLC performed experiments. DLC, ATP contributed to writing of the manuscript. All authors read and approved the manuscript.

## Acknowledgements

This publication and the underlying study have been made possible partly on the basis of the data that Hartwig Medical Foundation and the Center of Personalised Cancer Treatment (CPCT) have made available to the study.

## Notes

https://github.com/PapenfussLab/gridss

### Competing Interest Statement

The authors have declared no competing interest.

